# A Panel of Diverse *Pseudomonas aeruginosa* Clinical Isolates for Research and Development

**DOI:** 10.1101/2021.04.23.441230

**Authors:** Francois Lebreton, Erik Snesrud, Lindsey Hall, Emma Mills, Madeline Galac, Jason Stam, Ana Ong, Rosslyn Maybank, Yoon I. Kwak, Sheila Johnson, Michael Julius, Brett Swierczewski, Paige E. Waterman, Mary Hinkle, Anthony Jones, Emil Lesho, Jason W. Bennett, Patrick McGann

**Affiliations:** Multidrug-Resistant Organism Repository and Surveillance Network (MRSN), Walter Reed Army Institute of Research, Silver Spring, Maryland, USA; Bacterial Disease Branch, Walter Reed Army Institute of Research, Silver Spring, Maryland, USA; Office of Science & Technology Policy, Washington DC, USA; Infectious Diseases Unit, Rochester General Hospital, Rochester New York, USA; Department of Virology, Armed Forces Research Institute of Medical Sciences (AFRIMS), Bangkok, Thailand

**Author notes:** Francois Lebreton and Erik Snesrud contributed equally to this work. Corresponding Author, Patrick Mc Gann, PhD, Walter Reed Army Institute of Research, 503 Robert Grant Avenue, 2A36, Silver Spring, MD 20910, USA.

**Keywords:** *Pseudomonas aeruginosa*, antimicrobial resistance, whole-genome sequencing

## Abstract

*Pseudomonas aeruginosa* is a leading cause of community-acquired and hospital-acquired infections. Successful treatment is hampered by its remarkable ability to rapidly develop resistance to antimicrobial agents, mostly through mutation. In response, the World Health Organization listed carbapenem-resistant *P. aeruginosa* as a Priority 1 (Critical) pathogen for research and development of new treatments. A key resource in developing effective countermeasures is access to diverse and clinically relevant strains for testing. Herein we describe a panel of 100 diverse *P. aeruginosa* strains to support this endeavor.

Whole genome sequencing was performed on 3,785 *P. aeruginosa* housed in our repository. Isolates were cultured from clinical samples collected from healthcare facilities around the world between 2003 and 2017. Core-genome multi-locus sequence typing and high-resolution SNP-based phylogenetic analyses were used to select a panel of 100 strains that captured the genetic diversity of this collection. Comprehensive antibiotic susceptibility testing was also performed using 14 clinically relevant antibiotics.

This 100-strain diversity panel contained representative strains from 91 different sequence-types, including genetically distinct isolates from major epidemic clones ST-111, ST-235, ST-244, and ST-253. Seventy-one distinct antibiotic susceptibility profiles were identified ranging from pan-sensitive to pan-resistant. Known resistance alleles as well as the most prevalent mutations underlying the antibiotic susceptibilities were characterized for all isolates.

This panel provides a diverse and comprehensive set of *P. aeruginosa* strains for use in developing solutions to antibiotic resistance. The isolates, and all available meta-data including genome sequences, are available to industry, academic institutions, federal and other laboratories at no additional cost.

**Importance:** *Pseudomonas aeruginosa* is one of the most important human pathogens and a leading target in the development of new drugs and therapeutics. This species displays a remarkable level of diversity and any potential therapeutic must contend with this characteristic to ensure it retains efficacy across different strains. To date, only limited panels of *P. aeruginosa* are available for testing, and none have been designed to capture the genetic diversity of the species. The panel described herein has been designed to address this shortcoming by providing a set of 100 distinct strains that greatly captures the diversity of the species. This panel will be of significant value to research groups working on this important pathogen, both for research purposes and in the development of new diagnostics and countermeasures.

## Introduction

*Pseudomonas aeruginosa* is a ubiquitous organism whose genetic and metabolic versatility has enabled it to survive and thrive in a diverse range of environments (1). This adroit adaptability has allowed *P. aeruginosa* to emerge as one of the most successful opportunistic pathogens, as evidenced by its inclusion in the clinically important ESKAPE (*Enterococcus faecium, Staphylococcus aureus, Klebsiella pneumoniae, Acinetobacter baumannii, Pseudomonas aeruginosa*, and *Enterobacter* spp.) coterie of human pathogens (2). In humans, *P. aeruginosa* is an important source of both community and hospital acquired infections, ranging from skin and soft tissue infections to pneumonias and complicated bloodstream infections (3, 4). In particular, *P. aeruginosa* is a signature pathogen of patients with cystic fibrosis and a significant source of infection in burn wounds, resulting in substantial morbidity and mortality (4, 5).

The population structure of *P. aeruginosa* is as complex as it is varied and significant overlap between environmental and clinical isolates has been observed (6). More recent studies using whole genome sequencing (WGS) have further refined this structure, with the species being segregated into two major clades (hereafter referred to as Group A and Group B as per the nomenclature used by Ozer *et al*. (7)) and up to three smaller clades (7–11). Group A and B are most often associated with human infections and there is evidence that recombination between the two groups is limited, suggesting infrequent ecological overlap (7). Genetically, the two clades can be distinguished to a high degree of accuracy based on the presence of one of two genes encoding the major effector proteins secreted by the *P. aeruginosa* type III secretion system, *exoS* and *exoU*. Specifically, in a recent study of 739 *P. aeruginosa* from diverse environments, 98% of isolates from Group A carried *exoS* but not *exoU*, whereas 95% of Group B isolates carried *exoU* but not *exoS* (7). While isolates from both groups cause human infections, those carrying *exoU* have long been associated with more severe outcomes and higher mortality rates (12–15). The smaller clades range from one to three groups depending on the study (6–8), with the largest study defining 3 groups: 3, 4 and 5 (8). Among them the unusual PA7 clade (16) was consistently identified. This clade is a taxonomic outlier that groups isolates with seemingly reduced virulence compared to Group A and B, but increased biofilm formation and antibiotic resistance (17).

A major contributor to the success of *P. aeruginosa* has been its remarkable ability to resist the action of antibiotics (18, 19). The organism can employ a wide range of intrinsic mechanisms to resist these agents, including porin loss through mutation, hyperproduction of intrinsic AmpC enzymes and efflux pumps, and modification of antibiotic targets (for a recent review, see (19)). The challenges posed by these formidable intrinsic mechanisms have been further compounded by the emergence of multi-drug resistant and extensively-drug resistant (MDR/XDR) strains carrying a variety of transferable antibiotic resistance genes (ARG), including those encoding extended-spectrum β-lactamases (ESBLs), carbapenemases, and 16S rRNA methyltransferases (20–22). Contrasting with the high genetic diversity within the antibiotic susceptible population (21), MDR and XDR isolates have been closely associated with a small number of globally distributed clones (for a review see (21)). The most prevalent of these “high-risk” clones is ST-235 (the founding member of clonal complex 235), which appears to have arisen in Europe during the early 1980s, shortly after the introduction of fluoroquinolones for treating *Pseudomonas* infections (23). The MDR and XDR phenotypes exhibited by ST-235 and other “high-risk” clones, such as ST-111 and ST-175 (21), are a combination of both mutation-driven resistance and acquisition of transferable ARG, particularly extended-spectrum β-lactamases and carbapenemases (20). Of particular concern, these “high-risk” clones are increasingly being encountered worldwide, making control of these challenging pathogens even more difficult (21, 24, 25).

Though antimicrobial drug discovery has languished for the past 40 years, the emerging antibiotic resistance crisis has seen renewed interest by governmental, academic, and industrial organizations (26–28). In 2017, the World Health Organization (WHO) listed carbapenem-resistant *P. aeruginosa* as a Priority 1 (Critical) pathogen for research and development of new treatments (29). In addition, the United States Center for Disease Control (CDC) listed MDR *P. aeruginosa* as a serious public health threat that requires “prompt and sustained action”(30). Recent advances in genomic, proteomic and bioinformatic technologies have opened new avenues for developing new antimicrobial agents, including new strategies to combat *P. aeruginosa* (31). An important consideration when developing strategies to combat *P. aeruginosa* is determining the most appropriate strain to use (31). While the reference strains PA01 and PA14 are the most commonly used research strains worldwide, they are not “high-risk” clones and do not provide a comprehensive representation of the species.

Here, akin to our previous effort with *A. baumannii* (32), the construction of a reference panel of 100 strains that captures the genetic diversity of clinical *P. aeruginosa* and maximizes phylogenetic distance and pan-genome diversity is described. The panel was designed based on the whole genome sequencing of 3,785 *P. aeruginosa* housed at the Multi-drug resistant organism Repository and Surveillance Network (MRSN), which have been collected over the past 11 years from around the world. The panel has diverse antibiotic susceptibility phenotypes, ranging from pan-susceptible to pan-resistant, due to both intrinsic and acquired resistance mechanisms. This standardized resource will prove valuable for those interested in studying this important pathogen and will aid the design and development of novel antimicrobials and diagnostics.

## Methods

### *Pseudomonas aeruginosa* Repository

The MRSN collects and analyzes clinically-relevant multi-drug resistant organisms (MDROs) across the Military Healthcare System (MHS) (33) and around the world in collaboration with the U.S. Department of Defense’s (DoD) Global Emerging Infections Surveillance (GEIS) Branch. All samples are housed in a central repository that currently contains over 85,000 isolates, including 3,785 *P. aeruginosa* that were cultured from 2,078 patients between 2003 and 2017 (**Figure S1**). The majority (84.1%) were collected from patients in the U.S., including Alaska and Hawaii, though strains from South America (4.4%), the Middle East (4%), Europe (3%), Africa (2.4%) and Asia (2.1%) are also represented. The isolates were cultured from a wide diversity of clinical specimens, including respiratory (37.9%), urine (27.4%), wounds (18.7%), surveillance swabs (5%), blood (4%), sterile fluid (2%), and the environment (1%). Isolate source was unavailable for 4.7% of the strains.

### Antibiotic Susceptibility Testing

Antibiotic susceptibility testing (AST) was performed in the MRSN’s College of American Pathologists (CAP) accredited clinical lab using the Vitek 2 with the AST-95 and AST-XN09 cards (bioMerieux, NC, USA). Fourteen antibiotics were tested, which were separated into seven different categories based on the *P. aeruginosa* antimicrobial categories outlined by Magiorakos *et al*. (34) with the following exceptions: the phosphonic acid and polymixin categories were replaced with a new category encompassing the β-lactam/β-lactamase inhibitor agents, ceftazidime-avibactam and ceftolozane-tazobactam. Susceptibility to these agents was used to classify the isolates as susceptible (susceptible to all 14 agents), multi-drug resistant (MDR, non-susceptible to ≥1 agent in ≥ 3 antimicrobial categories), extensively drug-resistant (XDR, non-susceptible to ≥1 agent in all but ≤2 categories), pandrug-resistant (PDR, non-susceptible to all 14 agents), or non-MDR (non-susceptible to 1 or 2 categories only).

### Whole Genome Sequencing and analysis

Isolates were sequenced on an Illumina MiSeq or NextSeq benchtop sequencer (Illumina, Inc., San Diego, CA) and analyzed as previously described (32), except that *in silico* multi-locus sequence typing (MLST) was performed at pubMLST (35) using the scheme developed by Curran *et al*. (36). Presence or absence of the *exoU* and *exoS* type III effector genes was determined as described by Ozer *et al*. (7). Briefly, BlastN searches of the *exoU* (reference nucleotide locus ID PA14_51530 in strain UCBPP-PA14) and *exoS* (locus ID PA3841 in strain PAO1) genes were performed using default parameters against the genomic sequences of each isolate in the diversity panel.

### Refinement of the *P. aeruginosa* repository

To reduce redundancy in the initial 3,785 isolate set, successive isolates after the first from the same patient that shared the same sequence type (ST) were removed unless the isolates were cultured from a different body site (e.g. blood versus urine) or were cultured >6 months apart. All isolates from the same patient with different STs were retained. This refinement resulted in a final panel of 2,661 isolates (from 2,085 patients) available for analysis.

### cgMLST analysis and initial panel selection

The draft genomes of the 2,661 isolates were uploaded and analyzed using SeqSphere+ software (Ridom, Germany) using the *P. aeruginosa* core genome multi-locus sequence typing (cgMLST) scheme developed by Stanton *et al*. (37). To be included in the analysis, isolates had to contain 90% of the 4,400 genes included in this cgMLST scheme. The resulting Minimum Spanning Tree (MST) was then used to select 310 strains that captured the diversity of the strain collection.

### Core Genome SNP and Accessory Genome Analysis

PanSeq (38) was run with a fragmentation size of 500 bp to find sequences with ≥95% identity in ≥90% of the isolates to generate the core genome single nucleotide polymorphism (SNP) alignment and accessory genome binary alignment for the selected diversity set of 310 isolates. The SNP-based phylogeny was built using RAxML (version 8.2.11) (39) from a 534 kb variable position alignment using the GTR GAMMA model and the rapid bootstrapping option for nucleotide sequences (100 replicates). Using this approach, 100 strains were selected to represent the final diversity panel.

Prior to phylogenetic analysis, the set of 100 isolates was re-sequenced and re-analyzed to confirm purity. Reads were checked for contamination at the species level with Kraken2 (version 2.0.8-beta) (40) using the standard database build and at the strain level using ConFindr (version 0.4.8) (41) with parameters bf=0.05 and q=30. ConFindr parameters were validated for *P. aeruginosa* using a small subset of isolates that were deemed not contaminated by ConFindr using the default settings (q=20). For this validation set, paired reads were randomly selected using seqtk (https://github.com/lh3/seqtk) to a normalized read total of 3 million reads. These uncontaminated isolates were then mixed *in silico* with another isolate from a different rMLST scheme at 0-10, 15, 20, 25, and 50 percent of the reads. Panel isolates with greater than 1% of their reads belonging to a different species as identified by Kraken2, or those with 3 or more single nucleotide variants (SNVs) as identified by ConFindr, were further purified in the laboratory and re-sequenced. These purified sequences were compared to the original genomes by phylogeny to ensure identity. Finally, phylogenies were generated as described above.

Specifically, the core genome alignment was 1.7Mb long and contained 122 kb variable positions. Genome annotations were performed using NCBI Prokaryotic Genome Annotation Pipeline (version 4.8) and were made available on GenBank (BioProject: PRJNA446057).

### Analysis of chromosomal mutation-associated antibiotic resistance

No comprehensive tool, with a well-curated database, currently exists to accurately predict the mutational resistome of *P. aeruginosa*. In an attempt to identity the most well-characterized mutations, the existing literature was examined to obtain a non-exhaustive list of most frequently observed 151 amino acid substitutions in a set of 55 *P. aeruginosa* chromosomal genes that have previously been shown to be involved in antibiotic resistance (19, 42–46). For each gene, the corresponding protein sequences from the 100 *P. aeruginosa* panel isolates were aligned using MUSCLE (47) and the presence of known variants (including novel substitutions at known sites) was individually recorded (**Fig. 3** and **Table S1**).

### Diversity Panel Availability

The final diversity panel has been deposited at BEI resources (https://www.beiresources.org/) and is currently available for research purposes under catalogue # NR-51829.

## Results

### Strain diversity and initial isolate selection

All 2,661 de-duplicated *P. aeruginosa* passed the 90% gene threshold stipulated by cgMLST and were used to generate a minimum spanning tree (MST) (**Figure 1A**). The isolates were separated to a high degree of accuracy into two major clades by the presence of the *exoS* or *exoU* genes, consistent with the findings of Ozer *et al*. (7). One minor clade, consisting of 40 distantly related isolates that belong to the PA7 taxonomic outlier (16), was readily apparent while a second smaller group, consisting of 15 genetically divergent isolates, was only identified using phylogenetic analysis (**Figure 1b**). Notably, all 40 isolates from the PA7 clade and the 15 isolates from the smallest clade lack both *exoS* and *exoU*.

**Figure 1:**
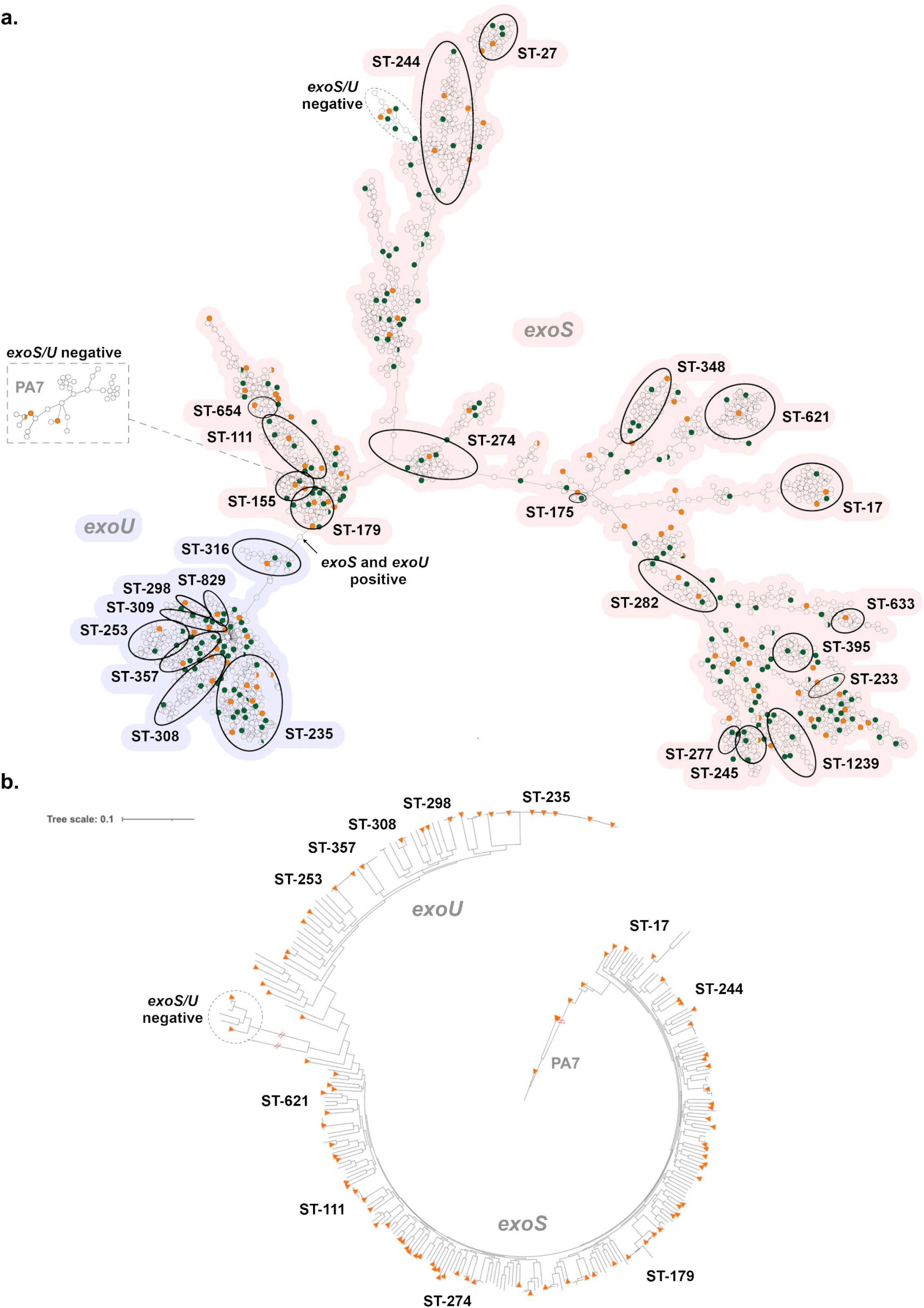
Genetic diversity of *P. aeruginosa* in the MRSN collection. **(A)** cgMLST minimum spanning tree of the 2,661 *P. aeruginosa* genomes used for initial panel selection. Isolates with an identical cgMLST allelic profile are represented by a single circle. For increased clarity, highly divergent isolates within the PA7 clade are displayed on a separate dendrogram wherein predicted root placement in the global population was indicated in image post-processing. The most prevalent and/or important ST (from traditional MLST typing) are indicated. The presence of *exoS* (red shading) or *exoU* (blue) is indicated for the two main clades. The absence of both alleles is noted for the PA7 and a second smaller clade. The 310 isolates initially selected to represent the breadth of diversity (see the text for details) are shown in green and orange with orange indicating the 100 strains in the final panel. **(B)** Core-genome SNP-based phylogenetic tree of 310 *P. aeruginosa*. The 100 isolates selected for the final panel are indicated with orange triangles. STs are indicated as well as the major clade designations.

Traditional MLST assigned 2,304 isolates to 407 known sequence types (STs) and the remaining 357 isolates to 274 novel STs. Seven of the top 10 most common STs in the repository have previously been detected in at least 3 countries (21) including ST-235 (9.7%; the most common clone), ST-244 (3.76%), ST-179 (2.86%), ST-253 (2.86%), ST-111 (2.82%), ST-17 (2.22%), and ST-274 (2.07%). Many other recognized clonal groups were also represented (**Figure 1a**).

### Selection of a non-redundant, genetically diverse panel of *P. aeruginosa*

The MST was used to select a subset of 310 strains that best represented the breadth and diversity of the overall strain collection (**Figure 1**). This subset encompassed 193 known STs as well as 38 strains with unique and novel STs whilst retaining the temporal and geographic distribution of the full strain collection. All 310 isolates were compared at the highest resolution using a SNP-based phylogenetic tree (**Figure 1b**). As illustrated, the overall population structure was retained (i.e. two large *exoU* and *exoS* clades as well as smaller PA7 and *exoU/S* negative clades) confirming the initial selection method using cgMLST. Ultimately, this phylogeny was used to select the final panel of 100 strains, chosen to maximize genetic diversity (**Figure 1b**).

The selected 100 strains were cultured from a wide range of clinical samples between 2003 and 2017, with the majority (n = 97) collected from hospitals across the U.S. (**Table S1**). Genetic diversity was high, with representatives from 91 different STs, including distinct isolates from the clinically important clones ST-111, ST-235, ST-244, and ST-253. All permutations of the *exoU* and *exoS* gene were represented; 72 isolates carried *exoS* (including MRSN 11278, which otherwise clustered with isolates from the *exoU* clade in the phylogeny), 21 carried *exoU*, two carried both, and five isolates lacked both genes (**Table S1**).

This substantial diversity is also reflected in the gene content, where just 3,200 core genes are shared by all 100 isolates, while >25,000 distinct genes are represented in the pan-genome (**Figure 2**). Furthermore, the size of the pan-genome never plateaus, highlighting the low redundancy and large amount of gene diversity in the panel.

**Figure 2:**
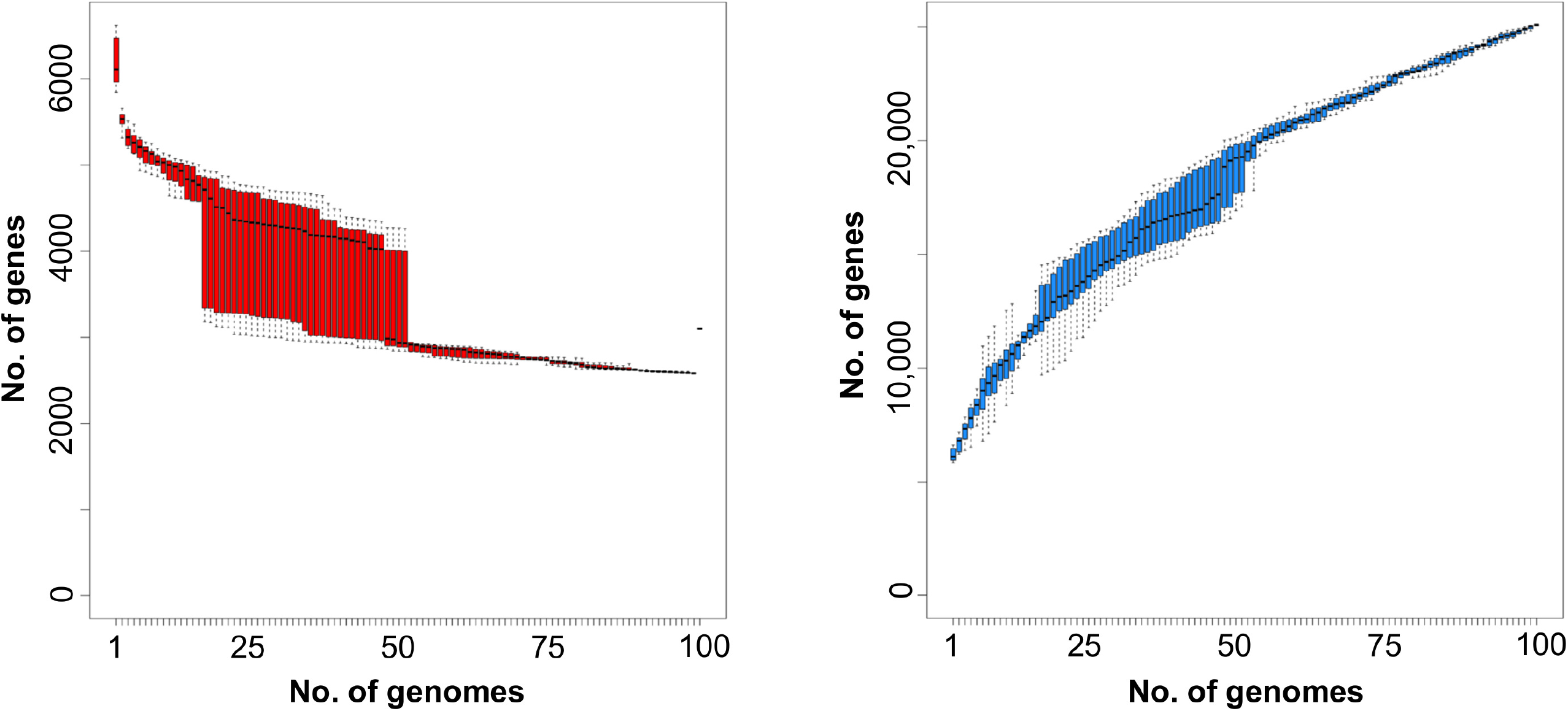
Core and pan genome of 100 diverse *P. aeruginosa*. Gene rarefaction (left, core genome) and accumulation (right, pan genome) curves are provided with the number of genes (y-axis) as a function of the number of genomes (x-axis), from a random sample (variation indicated as boxplot) of all genomes. Core genome is defined as the number of genes found in 99% of the genomes.

### Antibiotic susceptibility and antibiotic resistance mechanisms

Phenotypically, the selected isolates composing the diversity panel encompass a diverse range of susceptibilities; 20 isolates were susceptible to all 14 agents, 23 were non-MDR, 12 were MDR, 44 were XDR, and one isolate (MRSN 6220) was resistant to all 14 antibiotics (PDR) (**Figure 3, Table S1**). Overall, 71 distinct antibiotic susceptibility profiles were identified, including susceptible and non-susceptible strains across all seven categories. Notably, 64 isolates were non-susceptible to one or both carbapenems tested (imipenem and meropenem) and 19 and 12 isolates were non-susceptible to the newer β-lactam/β-lactamase combinations ceftazidime-avibactam and ceftolozane-tazobactam, respectively (**Figure 3**, **Table S1**).

**Figure 3:**
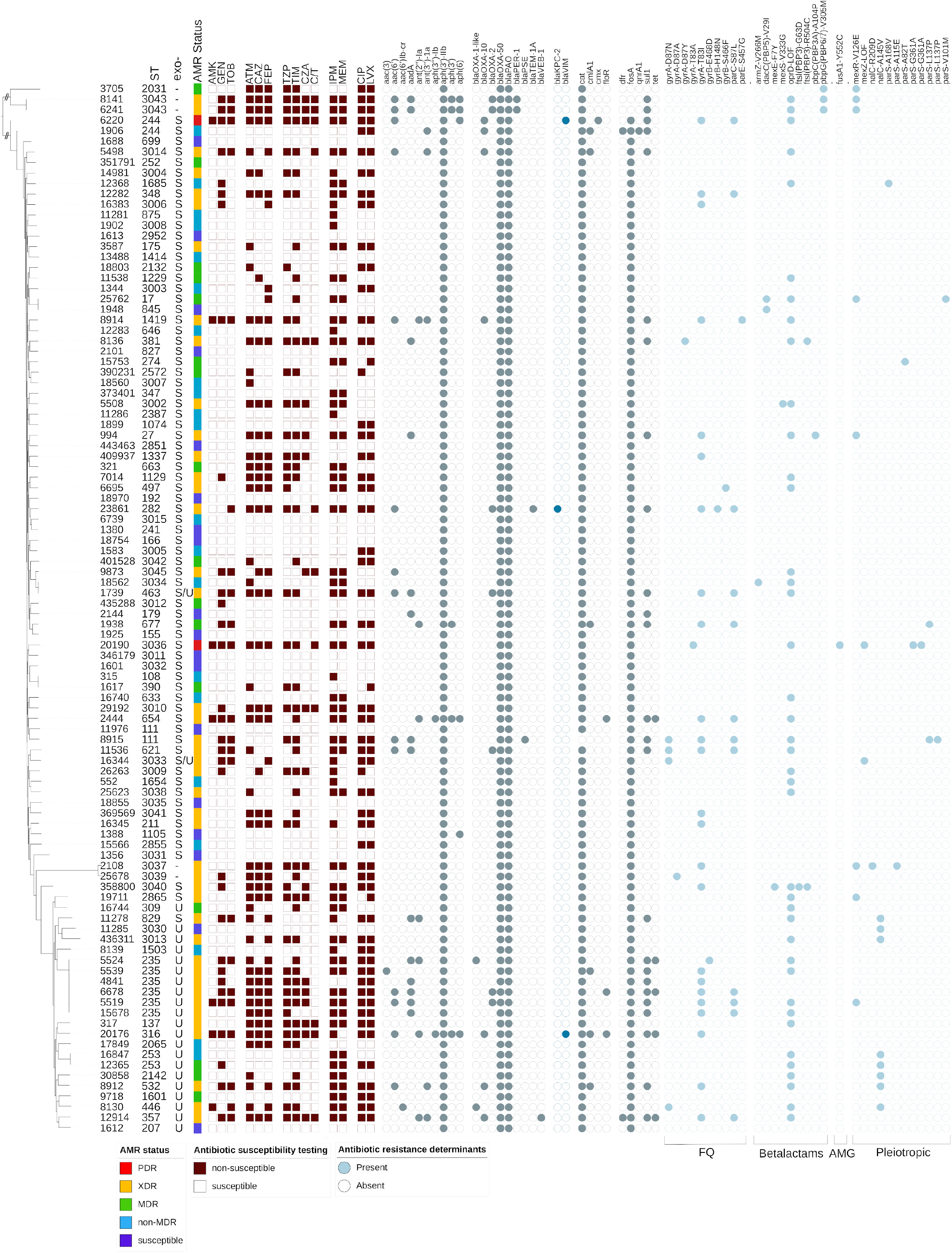
Characteristics of the *P. aeruginosa* diversity panel. Core genome SNP-based phylogenetic tree of the 100 strains in the final diversity panel. Strain sequence type (ST) is indicated. The assigned antimicrobial resistance (AMR) phenotype (see the text for details) is provided, and the dark red-brown squares indicate a result of non-susceptible (filled) or susceptible (open) to the tested antibiotic. The gray circles indicate the presence of a known resistance gene or mutation, with the blue-gray color highlighting those gene families of particular interest. For each isolate, the specific effector/genotype (*exoU+* or *exoS+*) of the type III secretion system are indicated.

Genetically, 88 antibiotic resistance genes (ARGs) were identified among the 100 isolates (**Table S1**), but this included 17 distinct alleles of the intrinsic *bla*_OXA-50_-like gene, 34 distinct alleles of the intrinsic *bla*_PDC_ genes and 15 distinct alleles encoding aminoglycoside modifying enzymes (AMEs) (**Table S1**). In addition to the two intrinsic β-lactamases, all isolates carry the kanamycin resistance gene *aph(3’)*-IIb and the chloramphenicol resistance gene *catB7*,while 99 of the 100 isolates carry the fosfomycin resistance gene *fosA*. Remarkably, 71 isolates carry only these five intrinsic genes but exhibit antibiotic susceptibility profiles ranging from pan-susceptible to XDR indicating a vast array of acquired resistances most likely caused by mutations (**Figure 3, Table S1**).

#### Aminoglycoside resistance

Only 6 isolates were non-susceptible (i.e. resistant or intermediate) to all three aminoglycosides tested. In addition, 14 were susceptible to amikacin but non-susceptible to gentamicin and tobramycin, 12 were resistant to gentamicin alone, 2 were resistant to tobramycin alone and the remaining 66 were susceptible to all three. Overall, resistance to aminoglycosides correlated well with the presence of AME genes. For example, specific alleles of aminoglycoside N-acetyltransferases (e.g. *aac(6)*-Ib conferring resistance to amikacin (low-level) and tobramycin) and aminoglycoside O-nucleotydyltransferases (e.g. *ant(2”)-Ia* conferring resistance to gentamicin and tobramycin) were identified in 21 non-susceptible isolates (**Figure 3, Table S1**). Similarly, isolates susceptible to all aminoglycosides lacked these enzymes.

Thirteen isolates displayed variable aminoglycoside resistance in the absence of known AMEs, including 11 non-susceptible to gentamicin, 1 non-susceptible to tobramycin, and 1 non-susceptible to all three (MRSN 20190). In the absence of acquired aminoglycoside-modifying enzymes, aminoglycoside resistance in *P. aeruginosa* has been linked to multiple mechanisms, but mutational overexpression of the efflux pump system MexXY-OprM is the most common (19, 48). In this study, a non-exhaustive search for mutations in *mexZ* and/or *parRS* that have previously been linked to MexXY-OprM overexpression and aminoglycoside resistance was performed. A loss-of-function mutation in MexZ was identified in two gentamicin and tobramycin non-susceptible strains (MRSN 16344 and MRSN 20190), and a *parS* mutation was identified in gentamicin non-susceptible MRSN 12368, but the mechanism(s) underlying aminoglycoside resistance in the remaining 10 strains remains to be determined.

#### Carbapenem resistance

Of high clinical importance, 48 isolates were non-susceptible to both carbapenems (imipenem and meropenem) and 14 were non-susceptible to imipenem alone (**Figure 3, Table S1**). Notably, just 3 isolates carried an acquired carbapenemase (*bla*_VIM-11_ in MRSN 20176, *bla*_VIM-6_ in MRSN 6220 and *bla*_KPC-2_ in MRSN 23861) and one, MRSN 9873, carried a gene that shared 92% nucleotide identity over 98% of the gene length with *bla*_HMB-1_, a rare metallo-β-lactamase originally described in a clinical strain of *P. aeruginosa* in 2017 (49). Besides gene acquisition, a total of 40 strains carried a loss-of-function mutation in the outer membrane protein OprD, by far the most common mechanism for carbapenem resistance in this species (50) and a known modulator of colonization and virulence in a mouse model (51). A known mutation in ParS and loss-of-function mutation in MexZ were identified as the possible cause of carbapenem resistance in an additional 4 isolates while the mechanism(s) of carbapenem resistance remains undetermined for sixteen isolates (**Figure 3, Table S1**).

#### Fluoroquinolone resistance

From the diversity panel, 53 strains were non-susceptible to both fluoroquinolones (FQ) (ciprofloxacin and levofloxacin), 2 were non-susceptible to just ciprofloxacin, 3 were non-susceptible to levofloxacin only, and the remaining 42 were susceptible to both (**Figure 3, Table S1**). In just two isolates, MRSN 1906 and MRSN 8130, FQ resistance could be attributed to the acquired FQ resistance genes *qnrA* and *aac(6)-Ib-cr5*, respectively. This is consistent with the observation that high-level FQ resistance in *P. aeruginosa* is primarily driven by mutation, particularly mutations in the quinolone resistance-determining regions (QRDR) of DNA Gyrase (*gyrA* and *gyrB*) and topisomerase IV (*parC* and/or *parE*) and/or overexpression of the MexEF-OprN or MexCD-OprJ efflux pumps (19, 52). A non-exhaustive search for mutations known to cause FQ resistance in *P. aeruginosa* identified strains with mutations in *gyrA* (n=29), *gyrB* (=3), *parC* (n=12), and *mexR* (n=6). However, the mechanism(s) for the observed resistance to ciprofloxacin and/or levofloxacin in an additional 22 strains remains to be determined (**Figure 3, Table S1**).

## Discussion

The World Health Organization (WHO) and the United States Centers for Disease Control (CDC) have classified MDR and carbapenem-resistant *Pseudomonas aeruginosa* as one of the most serious antibiotic resistant threats (29, 30). These classifications highlight the clinical importance of *P. aeruginosa* and the increasing difficulty faced by clinicians in treating infections by this organism. Fortunately, this recognition has spurred renewed interest in developing effective countermeasures, including vaccines, antimicrobial peptides, antibodies, virulence inhibitors, bacteriophage, and novel antimicrobials (53, 54).

A key component in evaluating any new therapeutic or diagnostic is access to a diverse set of strains that capture the genetic diversity within a species. Access to a diverse source of strains can be difficult and as a result there is very little consistency in the strains used across studies. The urgency of this requirement has been recognized at the highest level of the U.S. Government, with the U.S. National Action Plan (NAP) for Combating Antibiotic-Resistant Bacteria (CARB) indicating the need for a “a specimen repository to facilitate development and evaluation of diagnostic tests and treatments”. Notably, the recently released CARB 2020-2025 document directs Federal Agencies to “continue expanding and improving access to specimen and data repositories for research and innovation” (55).

Since the publication of the first CARB national strategy and action plan in 2015, U.S. Federal agencies have strived to develop appropriate specimen repositories. For *P. aeruginosa*,the U.S. CDC and Food and Drug Administration (FDA) have developed a useful panel of 55 *P. aeruginosa* isolates that were chosen to represent a diversity of AST results for drugs that are used to treat infections (https://wwwn.cdc.gov/ARIsolateBank/Panel/PanelDetail?ID=12; last accessed April 20^th^, 2021). Notably, the strains carry a variety of antibiotic-resistant genes, including the potent carbapenemases *bla*_IMP_, *bla*_KPC_, *bla*_NDM_, and *bla*_VIM_. While this is a valuable resource for studying strains with specific antibiotic resistance patterns, there is no data on the phylogeny of the strains and how representative they are of the overall genetic diversity within this species. In 2013, De Soyza *et al*. created a panel of 43 diverse *P. aeruginosa* strains selected through consensus by a panel of clinical and research experts (56). However, this panel lacks comprehensive genome information, provides no antibiotic susceptibility data, and was generated from a limited dataset of just 955 genotyped isolates. In the intervening 8 years, new strains of *P. aeruginosa* with epidemic potential have emerged (21) and there is an urgent need to develop a more comprehensive panel of isolates for research and development efforts.

In an effort to address these limitations and provide a new panel that is complementary to these existing sets, the large repository of *P. aeruginosa* clinical isolates present in the MRSN was leveraged. These strains have been collected over the past 12 years from clinical samples around the world (**Figure S1**) and bolstered by historical strains stored in U.S. Military Treatment Facility (MTF) archives. Though the majority of strains were collected in the U.S., it is notable that those from other continents display a similar distribution, a likely consequence of the successful spread of high-risk clones across the globe (**Figure S1**). In addition to providing representatives of the major high-risk clones, including ST-111, ST-235, and ST-175, the panel also includes emerging clones of concern such as ST-244, ST-357, and ST-654 (20) as well as sporadic clones that capture the genetic diversity across all major clades (including taxonomic outlier PA07) previously identified for this species (6–8, 16).

Though the panel was designed toward maximizing genetic diversity, the selected strains of *P. aeruginosa* strains encompass a diverse range of antibiotic susceptibility patterns from pan-sensitive to pan-resistant when tested against 14 relevant antibiotics (**Table S1**). Unlike the MRSN’s panel of diverse *A. baumannii* isolates, only a minor correlation between antibiotic susceptibility and the presence of transferable ARG in *P. aeruginosa* was observed. This is a hallmark of *P. aeruginosa* and reflects the outsized role played by point mutations in driving antibiotic resistance in this species (52). Although non-exhaustive, literature reviews (18, 19, 22, 42, 44, 46, 48, 50, 52) were performed to identify, within the panel isolates, an array of mutations known to contribute to aminoglycoside, carbapenem, and fluoroquinolone resistance. As a result, only 10-22% of the strains are phenotypically resistant due to unresolved genetic determinants. Finally, the panel includes strains with variable resistances to more contemporary agents, such as ceftazidime-avibactam and ceftolozane-tazobactam, which will be a valuable resource for investigating resistance mechanisms or developing the next generation of countermeasures. Altogether, this wide representation of resistance mechanisms (and combinations thereof) is the direct, desired consequence of the methodology used for selecting strains composing this panel.

In summary, an extensive collection of over 3,500 *P. aeruginosa* was utilized to design a novel panel of 100 diverse strains that: (i) encompass the genetic diversity of this species, (ii) capture both major global epidemic clones and sporadic strains and (iii) display diverse resistance mechanisms and antibiotic susceptibility profiles when tested against a panel of 14 relevant antibiotics. The expectation is that easy access to the strains combined with high-quality draft genomes will facilitate multiple avenues of research into this critical human pathogen.

## Acknowledgements

We acknowledge the participation of clinical laboratories across the Military Healthcare System, GEIS-affiliated overseas laboratories, and numerous collaborators for their contributions of bacteria to the MRSN *Pseudomonas aeruginosa* repository.

This study was funded by the U.S. Army Medical Command and the Defense Medical Research and Development Program. The manuscript has been reviewed by the Walter Reed Army Institute of Research. There is no objection to its presentation. The opinions or assertions contained herein are the private views of the authors and are not to be construed as official, or reflecting the views of the Department of the Army or the Department of Defense.

---

**Supplemental Figure 1. Geographic diversity of *P. aeruginosa* strains.** cgMLST Minimum Spanning Tree of the 3,785 *P. aeruginosa* used for initial panel selection color-coded by the continent where they were cultured. The presence/absence of the *exoU/exoS* genes are provided for context (See text). See Figure 1 legend for additional details.

